# Polymorphism within the mitochondrial genome of the ctenophore, *Pleurobrachia bachei* and its ongoing rapid evolution

**DOI:** 10.1101/366880

**Authors:** Andrea B. Kohn, Leonid L. Moroz

## Abstract

The mitochondrial genomes in ctenophores are among the most compact in the animal kingdom with multiple rearrangements and examples of gene loss. Here, by resequencing of the *Pleurobrachia bachei* mitochondrial genome, we show that the high level of polymorphism (>10%) in *Pleurobrachia* might contribute to the ongoing fast evolution of ctenophores including the presence of truncated versions of apparently canonical genes such as ***cox1***. Second, the codon interpretations in ctenophores, without robust proteomic data related to mitochondrial genes, is still a challenging issue, which is open for future experimental analyses.

---

Mitochondrial genomes from non-model/under-investigate organisms can be challenging to analyze, and ctenophores are one of the most difficult cases. Ctenophora or comb-jellies is a lineage of basal metazoans with relatively compact mitochondrial and nuclear genomes^1,2^. The initial and the most recent phylogenomic analyses of nuclear genomes and comparative analyses of 37 transcriptomes from representatives of this group indicated that Ctenophora is the sister lineage to the rest of extant animals^1,3,4^. However, it was also suggested that ctenophores might undergo an evolutionary bottleneck around the Permian time with a potential loss of their ancestral diversity and subsequent radiation and rapid evolution^4^. Mitochondrial genomes in ctenophores were affected by such complex evolutionary history and lost a significant fraction of the ancestral mitochondrial gene complement in Metazoa.

In 2011/2012 we showed that the mitochondrial genome of the ctenophore *Pleurobrachia bachei*^5^ was highly divergent and fast evolving compared to other animals. Simultaneously published analysis of the mitochondrial genome of *Mnemiopsis leidyi^6^* came to similar conclusions. It was also discovered that ctenophore genomes are the smallest of all metazoan genomes sequenced to date.

Recently three mitochondria genomes from the benthic ctenophores *Coeloplana loyai, Coeloplana yulianicorum* and *Vallicula multiformis*^7^ were sequenced with a discovery of even the smallest animal mitochondrial genome (9,961 bases in *Vallicula multiformis*^7^). The data presented in this study further confirm that ctenophore mitochondrial genomes are highly divergent. Here, we indicate that the same is true even within the same species from this clade.

Arafat *et al.* also suggested that the comparative data from the benthic Platyctenida infer a possible need for reannotation of the original *P. bachei* mitochondrial genome^7^. Although more detailed annotation is certainly desirable, here we show that the level of polymorphism in the *Pleurobrachia* might contribute to the ongoing rapid evolution of ctenophore genomes including the presence of truncated versions of apparently canonical genes. Second, the codon interpretations in ctenophores, without robust proteomic data related to mitochondrial genes is still a challenging issue, which opens future experimental analyses.

## The resequencing and reannotation of the *Pleurobrachia* mitochondrial genome

The recent comparative analyses of Platyctenida mt genomes^7^ suggested “*erroneous sequencing in the P. bachei cox1 gene”.* At the time of the original publication^5^, no other ctenophore mitochondrial sequences were available to compare with *Pleurobrachia*. Thus, we now investigated this point by looking at the more confounding issue of possible polymorphism. The *Pleurobrachia* mtDNA genome was PCR amplified, cloned and then sequenced from a single animal and several animals by Sanger technology. To date, Sanger sequencing is the gold standard for all sequencing.

Arafat *et al.^7^* proposed that the “region upstream to the *cox1* gene includes several poly-T, and thus, they suspect the introduction of a frame shift due to erroneous sequencing of the number of T in these homopolymers” in the *Pleurobrachia* mtDNA genome. To sequence and annotate *Pleurobrachia* mtDNA genome, we also analyzed the original 454-genome assembly, Illumina, and the Sanger sequencing to validate each other mutually, and none of these technologies were intended to be used exclusively.

In contrast to next-generation sequencing, homopolymers are well-captured in Sanger sequencing. The recently implemented reannotation^7^ used only one *P. bachei* nuclear genome run, SRR1174875^1^ submitted to NCBI two years after the original *P. bachei* mtDNA genome paper was published, and concluded the presence of an extra T as the effect of erroneous sequencing.

The current overall *P. bachei* whole genome shotgun sequencing project BioProject: PRJNA213480, is comprised of fourteen nuclear genomic DNA sequencing projects that were assembled and deposited under GenBank: AVPN00000000.1^1^. Our *P. bachei* assembly contains one contig representing the mitochondrial genome and <u>does not</u> contain an extra T in the *cox1* gene.

Secondly, we decided to further examine the issue of polymorphism in the *Pleurobrachia* mtDNA genome by cloning and then sequencing this region of interest with Sanger technology from ten individual *P. bachei.* Of the ten individual animals analyzed six had the extra T or 60% (see **Fig.1a and b**). Thus, we can conclude that this deletion/insertion of a T is not erroneous sequencing as suggested^7^, but a reflection of high polymorphism within the species.

**Fig 1a.**
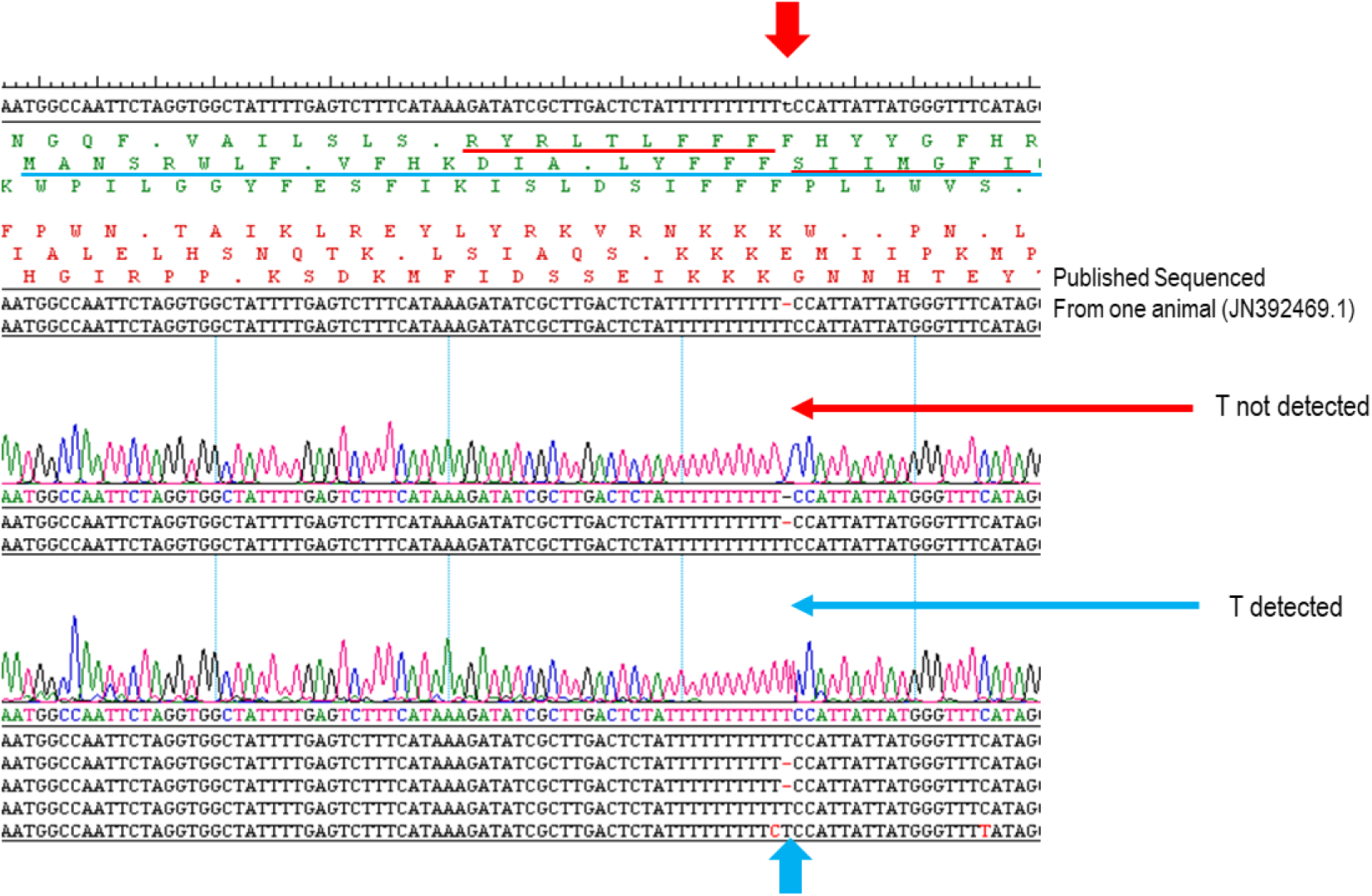
*Pleurobrachia cox1* sequence alignment from ten individual animals. Six contain the T-insertion and four do not. The red arrows indicate that the T-insertion is not detected as in the published *P. bachei* mitochondrial genome. The blue arrows specify individuals that had a T-insertion in the *cox1* gene. Possible frame shift is highlighted in the corresponding colors associate with the T insertion/deletion. The chromatograms are produced by Sanger sequencing technology and the most accurate tests in evaluating DNA sequencing (sequences were viewed in SeqMan, DNASTAR Inc, and corrections to the codon usage and protein translation were not possible in the SeqMan program). Only two example chromatograms are shown due to space.

**Fig 1b.**
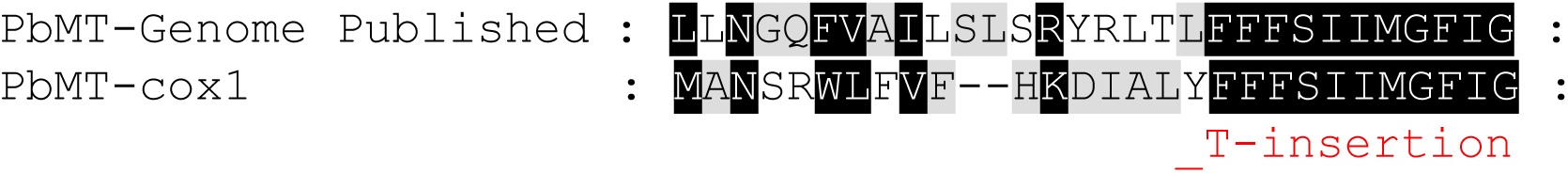
*P.bachei* predicted cox1 protein alignment. The red dash indicates the point of a T insertion and a frame-shift in the predicted coding sequence.

Next, we then checked available ctenophore transcriptomes to see if we could detect a deletion/insertion of a T at a similar position in the *cox1* gene. Both *Mnemiopsis leidyi* and *Vallicula multiformis* had sequences that detected a T-deletion, **Fig 2.** in a similar position as the *P. bachei cox1* gene.

**Fig 2.**
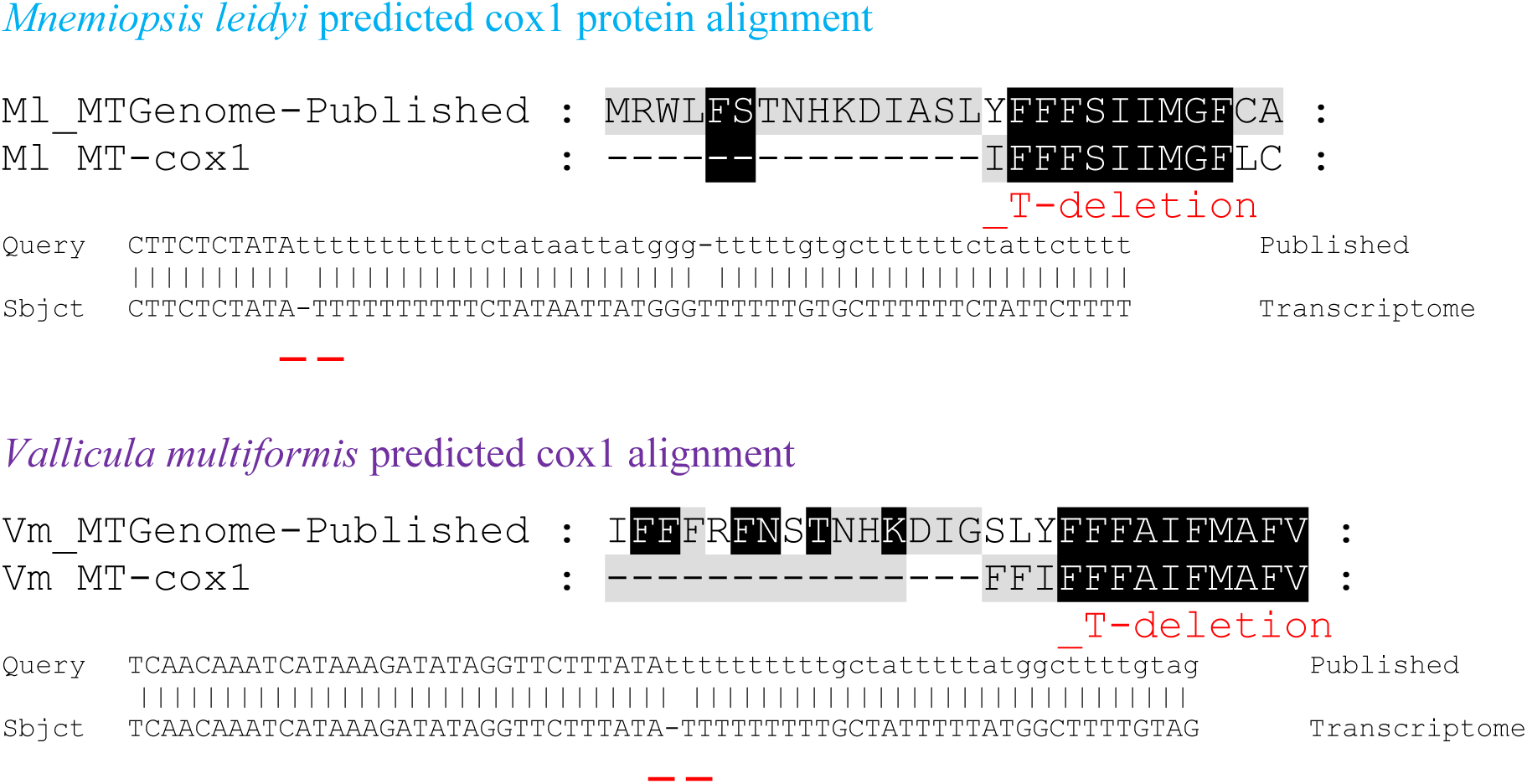
*Mnemiopsis and Vallicula* predicted *cox1* protein alignment. Transcritomes are publicly available BioProject: PRJNA213480 and https://neurobase.rc.ufl.edu/. The red dashs indicates the point of a T-deletion and a frame-shift in the predicted coding sequence.

These data on other ctenophore species also suggest high polymorphisms in the mitochondrial genomes within the clade (as it was correctly captured in the original submission, JN392469.1). We estimated the polymorphism of the *Pleurobrachia* mitochondria genome to be approximately 10% in the Friday Harbor populations (Pacific North West). These data confirm the need to sequencing multiple single animals collected in their natural habitats.

## Comments on the codon usage in ctenophore genomes

Considering the suggested rapid evolution of the ctenophore mitochondrial genomes, the correct interpretation of the codon usage is also challenging, and no mitochondrial proteomic data are available to date from this animal lineage. Arafat *et al.^7^* reannotation states “*P. bachei* the codon TGA, which codes for tryptophan in other ctenophores, is reassigned to serine as indicated by Pett and Lavrov^8^”.

TGA codon corresponds to UGA in RNA intermediates to protein synthesis. Earlier, we noted “In most metazoan mitochondrial genomes the UGA codon is a Tryptophan residue (Trp), however the UGA codon is undetermined in *P. bachei’s* mitochondrial genome. Based on alignments with other conserved proteins, UGA has the potential to encode different amino acids (Table 1S and Table 3S in Supplementary Material)” ^5^. Thus, the UGA codon was not robustly designated as a Tryptophan residue^5^.

In the computational re-analysis of previously published comparative mitochondrial sequences and novel genome data from ctenophores^1,2^, Pett and Lavrov in 2015^8^ stated: “the mt-genome from *Pl. bachei* (NC_016697) using GenDecoder^9^ showed that **38%** of TGA codons in highly conserved positions correspond to serine”. However, a 38% probability in our opinion doesn’t warrant making a definitive identification at this moment. We feel it is best to state the UGA codon is undetermined in *P. bachei’s* mitochondrial genome^5^, and experimentally test this hypothesis by proteomic approaches using more than one ctenophore species.

Finally, in the analyses of Platyctenida mitochondrial genomes^7^, it was indicated: “the true nad3 was not identified by us” and the authors proceed to state “the *nad3* gene shares the same translation frame as the downstream *nad4* gene and possesses an incomplete stop codon”. However, no phylogenetic analysis on Nad3 was presented. In essence, a prediction of putative proteins, with no unique start and no stop, is also challenging without proteomic data in such enigmatic animals as ctenophores.

## Conclusion

We conclude that the original annotation for the *Pleurobrachia* mitochondrial genome Genbank: JN392469.1 is consistent with the data available today, and the existing population data suggest both high level of polymorphism and the ongoing rapid evolution within this lineage. Nevertheless, as soon as more comparative/population data would be available, the reassignment of the codon usage in Ctenophora should be performed and further validated by experimental proteomic approaches.

## The Supplement

Tables from the original annotation of the mtDNA in *Pleurobrachia* (Genbank: JN392469.1)^5^.

| Gene | Position | Stand | Start | Stop | Length<br>(bp) | Coding<br>(aa) | UGA |
| --- | --- | --- | --- | --- | --- | --- | --- |
| *cox 1* | 1-1464 | + | ATG | TAA | 1464 | 484 | (4) G, T, A |
| *cox 2* | 1585-2166 | + | ATG | TAA | 582 | 193 | (1) A |
| tRNA-G<br>(ACC) | 2483-2544 | - |  |  | 62 |  |  |
| tRNA-P<br>(TGG) | 3220-3277 | + |  |  | 58 |  |  |
| *nad4L* | 3344-3523 | + | ATG | TAA | 180 | 60 | (1) I |
| *lrRNA* | 3524-3940 | + |  |  | 416 |  |  |
| *cox 3* | 3979-4719 | + | ATG | TAA | 741 | 246 | (3) S, T |
| *nad4* | 5590-6837 | + | ATG | TAA | 1248 | 415 | (2) G, L |
| *cob* | 7035-8081 | + | ATG | TAA | 1047 | 348 | (6) S, E, P S, G, L |
| *nad5* | 8101-9597 | + | ATG | TAA | 1497 | 498 | (11) I, G, T, S, M, P |
| *nad1* | 9588-10436 | + | ATT | TAA | 849 | 282 | (3) L, V |
| *nad3* | 10728-10928 | + | ATG | TAA | 210 | 70 | 0 |

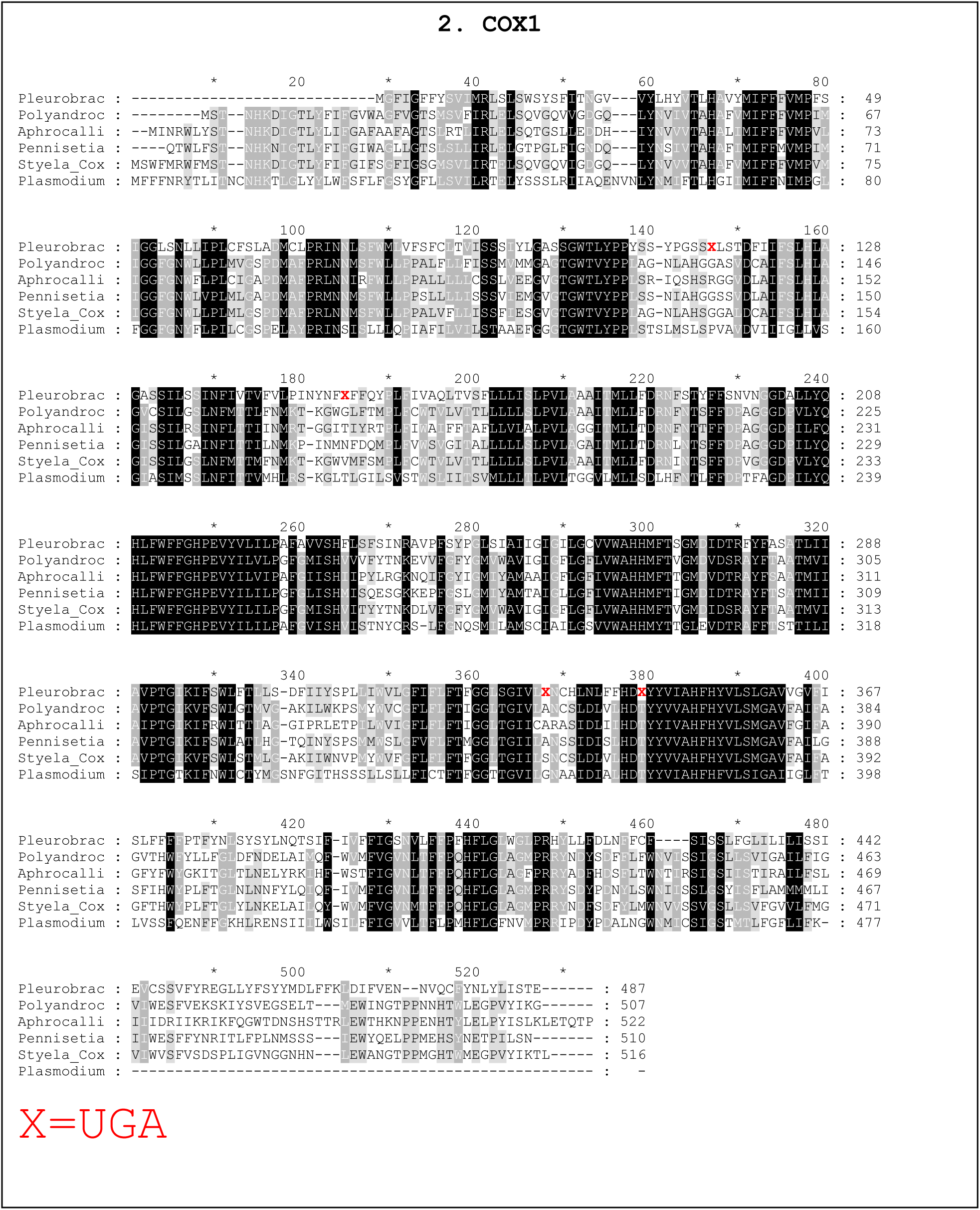

## Acknowledgments

This work was supported by the United States National Aeronautics and Space Administration (grant NASA-NNX13AJ31G), and the National Science Foundation (grants 1146575, 1557923, 1548121 and 1645219).

## References

1. Moroz, L. L. et al. The ctenophore genome and the evolutionary origins of neural systems. Nature 510, 109–114, doi:10.1038/nature13400 (2014).

2. Ryan, J. F. et al. The genome of the ctenophore *Mnemiopsis leidyi* and its implications for cell type evolution. Science 342, 1242592, doi:10.1126/science.1242592 (2013).

3. Whelan, N. V., Kocot, K. M., Moroz, L. L. & Halanych, K. M. Error, signal, and the placement of Ctenophora sister to all other animals. Proc Natl Acad Sci USA 112, 5773–5778, doi:10.1073/pnas.1503453112 (2015).

4. Whelan, N. V. et al. Ctenophore relationships and their placement as the sister group to all other animals. Nat Ecol Evol 1, 1737–1746, doi:10.1038/s41559-017-0331-3 (2017).

5. Kohn, A. B. et al. Rapid evolution of the compact and unusual mitochondrial genome in the ctenophore, *Pleurobrachia bachei*. Mol Phylogenet Evol 63, 203–207, doi:S1055-7903(11)00522-7 [pii] 10.1016/j.ympev.2011.12.009 (2012).

6. Pett, W. et al. Extreme mitochondrial evolution in the ctenophore *Mnemiopsis leidyi:* Insight from mtDNA and the nuclear genome. Mitochondrial DNA 22, 130–142, doi:10.3109/19401736.2011.624611 (2011).

7. Arafat, H., Alamaru, A., Gissi, C. & Huchon, D. Extensive mitochondrial gene rearrangements in Ctenophora: insights from benthic Platyctenida. BMC Evol Biol 18, 65, doi:10.1186/s12862-018-1186-1 (2018).

8. Pett, W. & Lavrov, D. V. Cytonuclear Interactions in the Evolution of Animal Mitochondrial tRNA Metabolism. Genome Biol Evol 7, 2089–2101, doi:10.1093/gbe/evv124 (2015).

9. Abascal, F., Zardoya, R. & Posada, D. GenDecoder: genetic code prediction for metazoan mitochondria. Nucleic Acids Res 34, W389–393, doi:10.1093/nar/gkl044 (2006).

